# Epigenetic clock and DNA methylation analysis of porcine models of aging and obesity

**DOI:** 10.1101/2020.09.29.319509

**Authors:** Kyle M. Schachtschneider, Lawrence B Schook, Jennifer J. Meudt, Dhanansayan Shanmuganayagam, Joseph A. Zoller, Amin Haghani, Caesar Z. Li, Joshua Zhang, Andrew Yang, Ken Raj, Steve Horvath

**Affiliations:** Department of Radiology, University of Illinois at Chicago, Chicago, IL, USA; Department of Biochemistry and Molecular Genetics, University of Illinois at Chicago, Chicago, IL, USA; National Center for Supercomputing Applications, University of Illinois at Urbana-Champaign, Urban, IL, USA; Department of Animal Sciences, University of Illinois at Urbana-Champaign, Urbana, IL, USA; Biomedical & Genomic Research Group, Department of Animal and Dairy Sciences, University of Wisconsin – Madison, Madison, Wisconsin, USA; Department of Surgery, University of Wisconsin School of Medicine and Public Health, Madison, Wisconsin, USA; Department of Biostatistics, Fielding School of Public Health, University of California, Los Angeles, Los Angeles, California, USA; Human Genetics, David Geffen School of Medicine, University of California, Los Angeles CA 90095, USA; Radiation Effects Department, Centre for Radiation, Chemical and Environmental Hazards, Public Health England, Chilton, Didcot, UK

**Keywords:** porcine, pig, minipigs, aging, development, epigenetic clock, DNA methylation

## Abstract

DNA-methylation profiles have been used successfully to develop highly accurate biomarkers of age, epigenetic clocks, for many species. Using a custom methylation array, we generated DNA methylation data from n=238 porcine tissues including blood, bladder, frontal cortex, kidney, liver and lung, from domestic pigs (*Sus scrofa domesticus*) and minipigs (Wisconsin Miniature Swine™). We present 4 epigenetic clocks for pigs that are distinguished by their compatibility with tissue type (pan-tissue and blood clock) and species (pig and human). Two dual-species human-pig pan-tissue clocks accurately measure chronological age and relative age, respectively. We also characterized CpGs that differ between minipigs and domestic pigs. Strikingly, several genes implicated by our epigenetic studies of minipig status overlap with genes (*ADCY3, TFAP2B, SKOR1*, and *GPR61*) implicated by *genetic* studies of body mass index in humans. In addition, CpGs with different levels of methylation between the two pig breeds were identified proximal to genes involved in blood LDL levels and cholesterol synthesis, of particular interest given the minipig’s increased susceptibility to cardiovascular disease compared to domestic pigs. Thus, inbred differences of domestic and minipigs may potentially help to identify biological mechanisms underlying weight gain and aging-associated diseases. Our porcine clocks are expected to be useful for elucidating the role of epigenetics in aging and obesity, and the testing of anti-aging interventions.

## Introduction

Pigs (*Sus scrofa*) are omnivores that last shared a common ancestor with humans between 79 and 97 million years ago ^1,2^. The domestication of pigs dates back to approximately 10,000 years, where they were bred with local wild boars across Eurasia ^3,4^. Since then, a wide variety of domestic and minipig breeds have been selectively bred for agricultural and biomedical purposes. While murine models have been traditionally used in biomedical research, there are added advantages to the use of porcine models in translational research. This include their comparable size, anatomy, physiology, immunology, metabolism, and genetics with humans ^5–7^. At the cellular levels, we have previously demonstrated similar genome-wide DNA methylation patterns between pigs and humans across a range of biomedically relevant tissues ^8,9^, further supporting the high relevance of pigs in modeling human disease and development. Indeed, these advantages have been recognized and porcine models are already being used in biomedical research ^10–19^. However, the large size and low propensity for atherosclerosis development in particular,^20^ limited the use domestic pigs as cardiovascular models. To overcome these issues, selectively bred and genetically modified minipigs have emerged over recent years as excellent models of hypercholesterolemia, atherosclerosis, metabolic syndrome, diabetes, and even cancer ^13,21–27^. Due to their smaller size, ease of handling, and genetic manipulability, minipigs are becoming increasingly important animal models for a wide range of human pathologies ^12^.

The study of any disease would be incomplete without an understanding of how age contributes to the malfunctioning of cells, tissues and organs. Despite this acknowledgement, the contribution of age to pathology has been largely unaddressed, not for the lack of will, but means. In the absence of an accurate way to quantify biological age, time (chronological age) is adopted as a surrogate that is manifestly unsatisfactory, as it remains unresponsive to biological fitness or frailty. The need for a measure of age that is based on biology is clear, and hints that this may be possible emerged when DNA methylation level was observed to change with advancing age. DNA methylation is an epigenetic modification that controls gene expression. The significance of its age-associated change was a subject of speculation until recently, when an array-based technology was developed to accurately measure its level on specific cytosine-phosphate-guanines (CpGs) in the genome. DNA methylation levels allow one to build age estimators (pan tissue clocks) that apply to most cells of the human body ^28^. The rate of epigenetic aging was observed to be associated with a wide range of human conditions including mortality risk, Down syndrome, HIV infection and obesity, ^29–36^ indicating that epigenetic age is a measure, at least to some degree, of biological age.

It is evident that the extrapolation of this epigenetic clock to other species, especially those such as pigs that are employed as disease models in biomedical research, will greatly facilitate research into the influence of age on pathology. This would also permit the quantitative testing of interventions that could potentially mitigate age effects on pathology. Towards this end, we aimed to develop epigenetic clocks that apply to both humans and pigs at the same time. To overcome the species barrier, we used a DNA methylation array platform (HorvathMammalMethylChip40) that encompasses CpGs flanked by DNA sequences that are conserved across different species of the mammalian class.

Here, we present highly accurate epigenetic clocks that apply to both humans and different pig breeds: the regular sized domestic pigs, Wisconsin Miniature Swine™, and a cross between domestic and Minnesota minipigs. The human-pig clock provides increases the probability that findings in pigs will translate to humans, and vice versa. We also characterized 1) age-related changes in the porcine methylome and 2) cytosines that differ between minpigs and regular sized pigs.

## Results

As detailed in the Methods, we used a custom methylation array (HorvathMammalMethylChip40) to generate DNA methylation data. In total, we analyzed 238 tissue samples mainly from blood (**Table 1**). Blood samples were obtained from three pig lines: a cross between the Large White and Landrace domestic pig breeds, the Wisconsin Miniature Swine minipig, and a cross between the Large White domestic breed (maternal line) and the Minnesota minipig (sire). From the latter pig line (i.e. the domestic minipig cross), methylation profiles were obtained from DNA isolated from bladder, brain (frontal cortex), kidney, liver, and lung tissue. Unsupervised hierarchical clustering revealed that the samples clustered by tissue type (**Supplementary Figure 1**). Random forest predictors were fitted to three different outcomes: 1) pig breed (*Sus scrofa domesticus* versus *Sus scrofa minusculus*), 2) tissue type, and 3) sex. The three classifiers exhibited perfect accuracy, with respective (out-of-bag) error rates of zero.

**Table 1.**
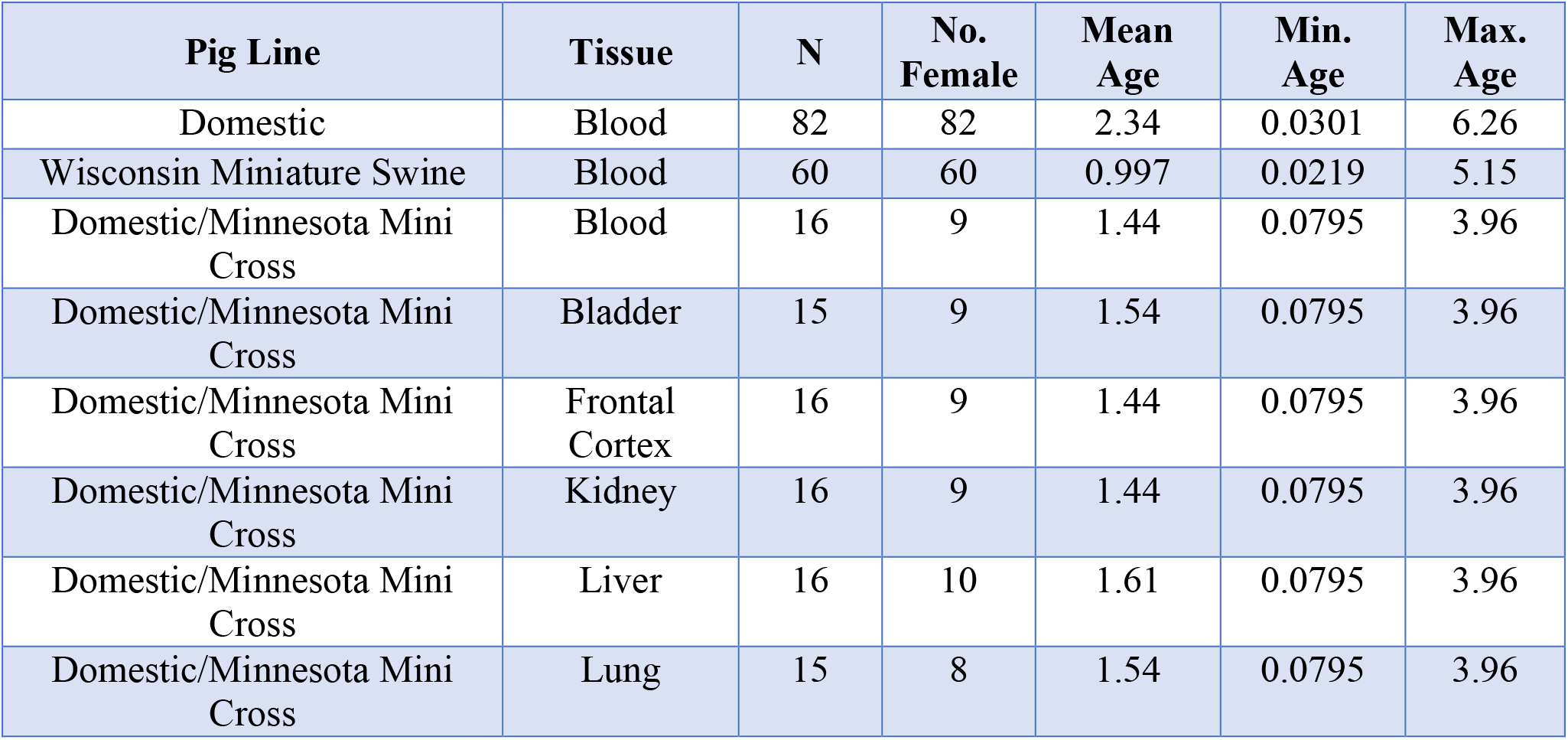
Description of the porcine methylation data. Pig lines. Tissue type, N=Total number of samples/arrays. Number of females. Age: mean, minimum and maximum.

| Pig Line | Tissue | N | No. Female | Mean Age | Min. Age | Max. Age |
| --- | --- | --- | --- | --- | --- | --- |
| Domestic | Blood | 82 | 82 | 2.34 | 0.0301 | 6.26 |
| Wisconsin Miniature Swine | Blood | 60 | 60 | 0.997 | 0.0219 | 5.15 |
| Domestic/Minnesota Mini Cross | Blood | 16 | 9 | 1.44 | 0.0795 | 3.96 |
| Domestic/Minnesota Mini Cross | Bladder | 15 | 9 | 1.54 | 0.0795 | 3.96 |
| Domestic/Minnesota Mini Cross | Frontal Cortex | 16 | 9 | 1.44 | 0.0795 | 3.96 |
| Domestic/Minnesota Mini Cross | Kidney | 16 | 9 | 1.44 | 0.0795 | 3.96 |
| Domestic/Minnesota Mini Cross | Liver | 16 | 10 | 1.61 | 0.0795 | 3.96 |
| Domestic/Minnesota Mini Cross | Lung | 15 | 8 | 1.54 | 0.0795 | 3.96 |

### Predictive Accuracy of the Epigenetic Clock

To arrive at unbiased estimates of the epigenetic clocks, we applied cross-validation analysis with the training data. For the development of the basic pig clock, this consisted of pig blood, bladder, frontal cortex, kidney, liver, and lung DNA methylation profiles. For the generation of human-pig clocks however, the training data was constituted by human and pig DNA methylation profiles. Cross-validation analysis reports unbiased estimates of the age correlation R (defined as Pearson correlation between the age estimate (DNAm age) and chronological age) as well as the median absolute error.

From these analyses, we developed three epigenetic clocks for pigs that vary with regards to two features: species and measure of age. The resulting two human-pig clocks mutually differ by way of age measurement. One estimates chronological ages of pigs and humans (in units of years) based on methylation profiles of 638 CpGs, while the other employs the methylation profiles of 542 CpGs to estimate relative age, which is the ratio of chronological age of an animal to the maximum lifespan of its species; with resulting values between 0 and 1. This relative age ratio is highly advantageous because it allows alignment and biologically meaningful comparison between species with very different lifespans such as pig and human, which cannot otherwise be afforded by direct comparison of their chronological ages.

As indicated by its name, the pure pig clock, constituted by 120 CpGs, is highly accurate in age estimation in all porcine tissues (R=0.97 and median error 0.22 years, **Figure 1A**). The pan-tissue clocks exhibit high age correlations in individual porcine tissues (R>0.90, **Supplementary Figure 1**). The human-pig clock for chronological age is highly accurate when DNA methylation profiles of both species are analyzed together (R=0.98, **Figure 1B**), and remains so when restricted to pig tissue samples (R=0.97, **Figure 1C**). Similarly, the human-pig clock for relative age exhibits high correlation regardless of whether the analysis is applied to samples from both species (R=0.98, **Figure 1D**) or only to pig samples (R=0.96, **Figure 1E**). The use of relative age circumvents the clustering of data points of pigs and humans to opposite parts of the curve, which is evident in Figure 1C. These highly accurate array of porcine clocks are readily useable with immediate effect in porcine models of diseases and conditions, and the human-pig clock of relative age is particularly exciting as it allows comparison between human and pigs based on their relative positions within the lifespans of both species.

**Figure 1.**
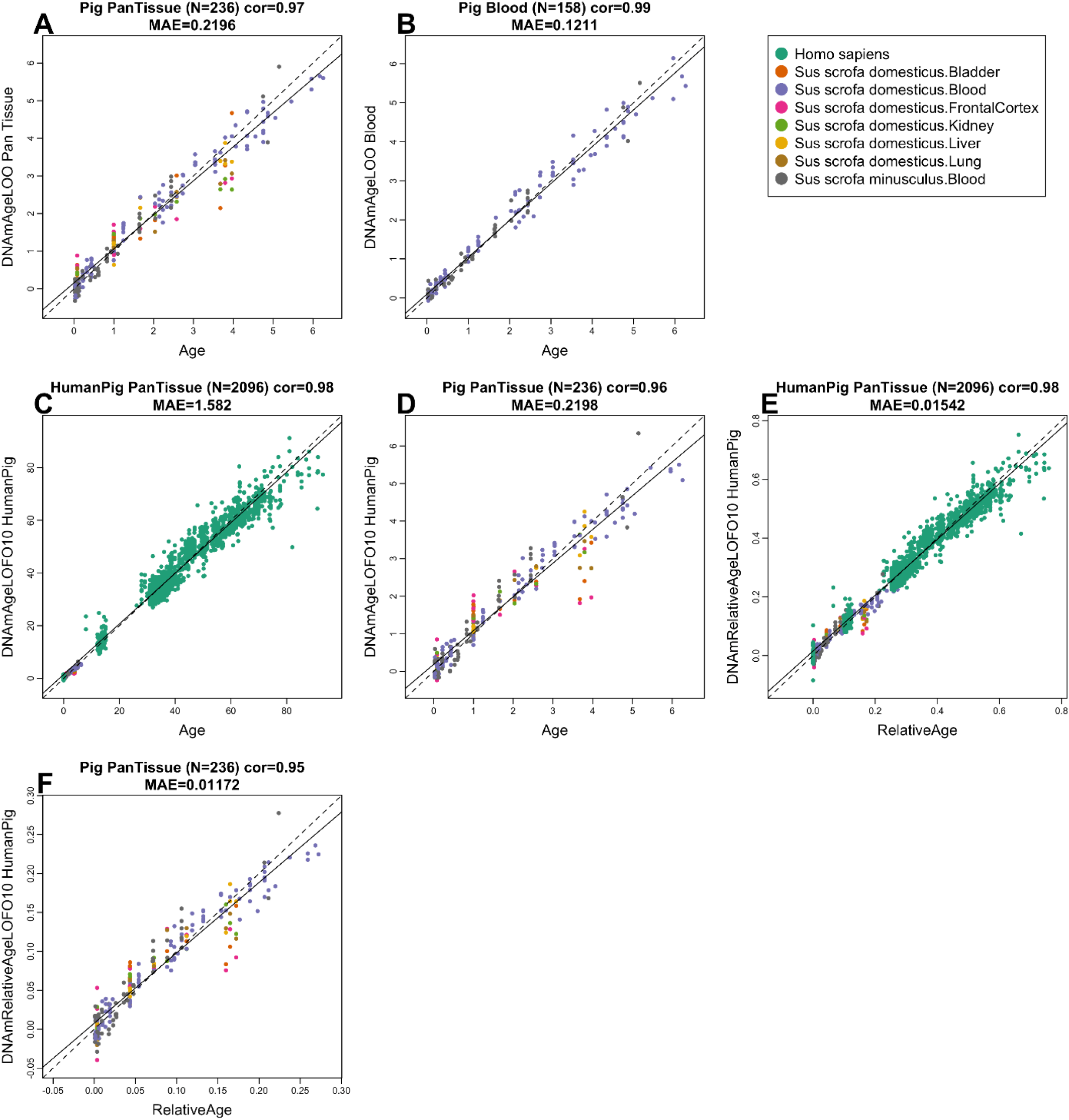
Cross-validation study of epigenetic clocks for pigs. We developed 4 epigenetic clocks for pigs: A) pan-tissue clock, B) blood clock, and human-pig clock for chronological age applied to C) both species and D) pigs only. Human-pig clock for relative age applied to E) both species and F) pigs only. Leave-one-sample-out (LOO) estimate (y-axis, in units of years) versus chronological age or relative age (x-axis). Relative age is defined as chronological age divided by the maximum age of the respective species. The linear regression of epigenetic age is indicated by a solid line while the diagonal line (y=x) is depicted by a dashed line.

### EWAS of chronological age

Although several hundred CpGs were used to construct the epigenetic clocks described above, these were merely a subset of all CpGs, which changed with advancing age. There are many more age-associated CpGs that are not used for the purpose of estimating porcine age, but are nevertheless very important when we seek to identify CpGs with methylation levels that are associated with age through epigenome-wide association studies (EWAS). In total, 34,540 probes from the HorvathMammalMethylChip40 are aligned to loci that are proximal to 5,209 genes in the *Sus_scrofa.Sscrofa11.1.100* genome assembly. Due to the high inter-species conservation of the probes on the array, findings from the pig methylation data can probably be extrapolated to humans and other mammalian species. EWAS of chronological age revealed clear tissue-specific DNAm change in pigs (**Figure 2A**). Age-associated CpGs in one tissue tend to be poorly conserved in another tissue (**Supplementary Figure 3**). However, the poor conservation and differences in p-value ranges in our analyzed tissue types may reflect the limited sample size in non-blood tissues. A nominal p-value < 10^-4^ was set as the cut-off for significance. The top age-associated CpGs and their proximal genes for the individual tissues are as follows: bladder, *ANO4* exon (z = 5.8); blood, *EN1* promoter (z = 26); frontal cortex, *FGF9* exon (z = 5.6); kidney, *TNRC6A* exon (z = −7.5); liver, *NRSA1* exon (z = 8.6); and lung, *UNC79* 5’UTR (z = 8.1). Despite poor conservation across tissues, there are nevertheless age-related CpGs that are common to all tissues, and these were identified through meta-analysis of these six tissue samples to be hypermethylation in *EN1* promoter (z = 18)*, HCN1* exon (z = 17), *UNC79* 5’UTR (z = 17), *LHFPL4* exon (z = 17), and *NR2F2* exon (z = 17). The upset plot analysis identified several CpGs with conserved DNAm aging in at least four pig tissues (**Figure 2B**). The most conserved DNAm aging pattern was hypermethylation of the *SP8* promoter in all tissues, with the exception of the brain. Genes whose expression are regulated by SP family transcriptional factors are essential for proper limb development. Aging-associated CpGs in different tissues were distributed in genic and intergenic regions that are defined relative to transcriptional start sites (**Figure 2C**). There is little enrichment of age-related CpGs in any genetic features across all tissues, with the exception of consistent hypermethylation in promoters and 5’UTRs. This result is consistent with a higher positive association of CpG island methylation with age than non-island CpGs in all tissues (**Figure 2D**). These features suggest that a substantial amount of age-associated CpGs are likely to impact gene expression.

**Figure 2.**
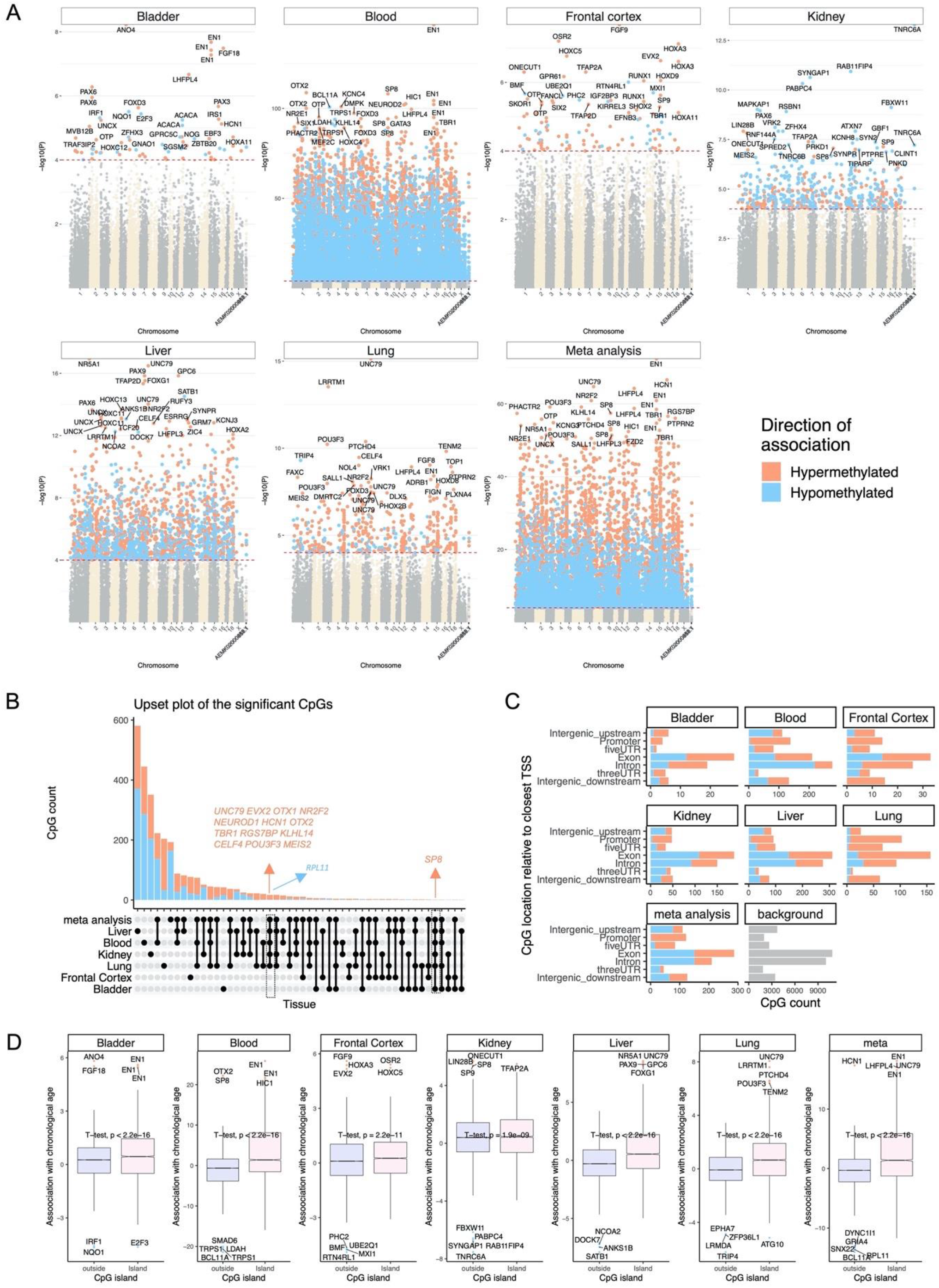
Epigenome-wide association (EWAS) of chronological age in porcine tissues. EWAS of age in bladder, blood, frontal cortex, kidney, liver, lung, and tissue meta-analysis of pigs (*Sus scrofa*). A) Manhattan plots of the EWAS of chronological age. The coordinates are estimated based on the alignment of the Mammalian array probes to the Sscrofa11.1.100 genome assembly. The direction of associations with p < 10-4 (red dotted line) is highlighted by red (hypermethylated) and blue (hypomethylated) colors. The top 30 CpGs were labeled by their neighboring genes. B) Upset plot representing the overlap of aging-associated CpGs based on meta-analysis or individual tissues. Neighboring genes of the overlapping CpGs are labeled in the figure. C) Location of top CpGs in each tissue relative to the closest transcriptional start site. Top CpGs were selected at p < 10-4 and further filtering based on z score of association with chronological age for up to 500 in a positive or negative direction. The number of selected CpGs: bladder, 73; blood, 1,000; frontal cortex, 123; kidney, 574; liver, 1,000; lung, 528; and tissue meta-analysis, 1,000. The grey color in the last panel represents the location of the 34,540 mammalian BeadChip array probes mapped to Sscrofa11.1.100 genome. D) Boxplot of DNAm aging for CpGs located within or outside CpG islands in porcine tissues. Labels indicate neighboring genes of the top 10 CpGs in each analysis.

To specifically explore the potential impact of age-related porcine CpG methylation on gene expression, we analyzed putative transcription factors binding sites for such methylation. From these we identified 20 transcription factor binding motifs that exhibit age-related methylation changes (**Figure 3A**). The corresponding transcription factors control the expression of genes which are involved in many different cellular activities. For example, hypomethylation in the SP1 motif in blood and cortex indicates greater access of the SP1 protein to some of its binding sites with increasing age. However, the outcome of this is difficult to predict as SP1 activates the transcription of many genes that are involved in diverse cellular processes ranging from cell growth, apoptosis, the immune response and chromatin remodeling. The challenge in predicting downstream events from single transcription motifs can be partly addressed by collective analysis of multiple transcription factor motifs. Such an analysis identified age-associated methylation changes for SMAD3, SP1, SP3, and E2F1 transcription factor binding motifs, which are implicated in telomerase regulation (p = 6E-7). We briefly mention that the effects of telomere length and telomerase activity in porcine tissues appears to be comparable with that of humans ^37^.

**Figure 3.**
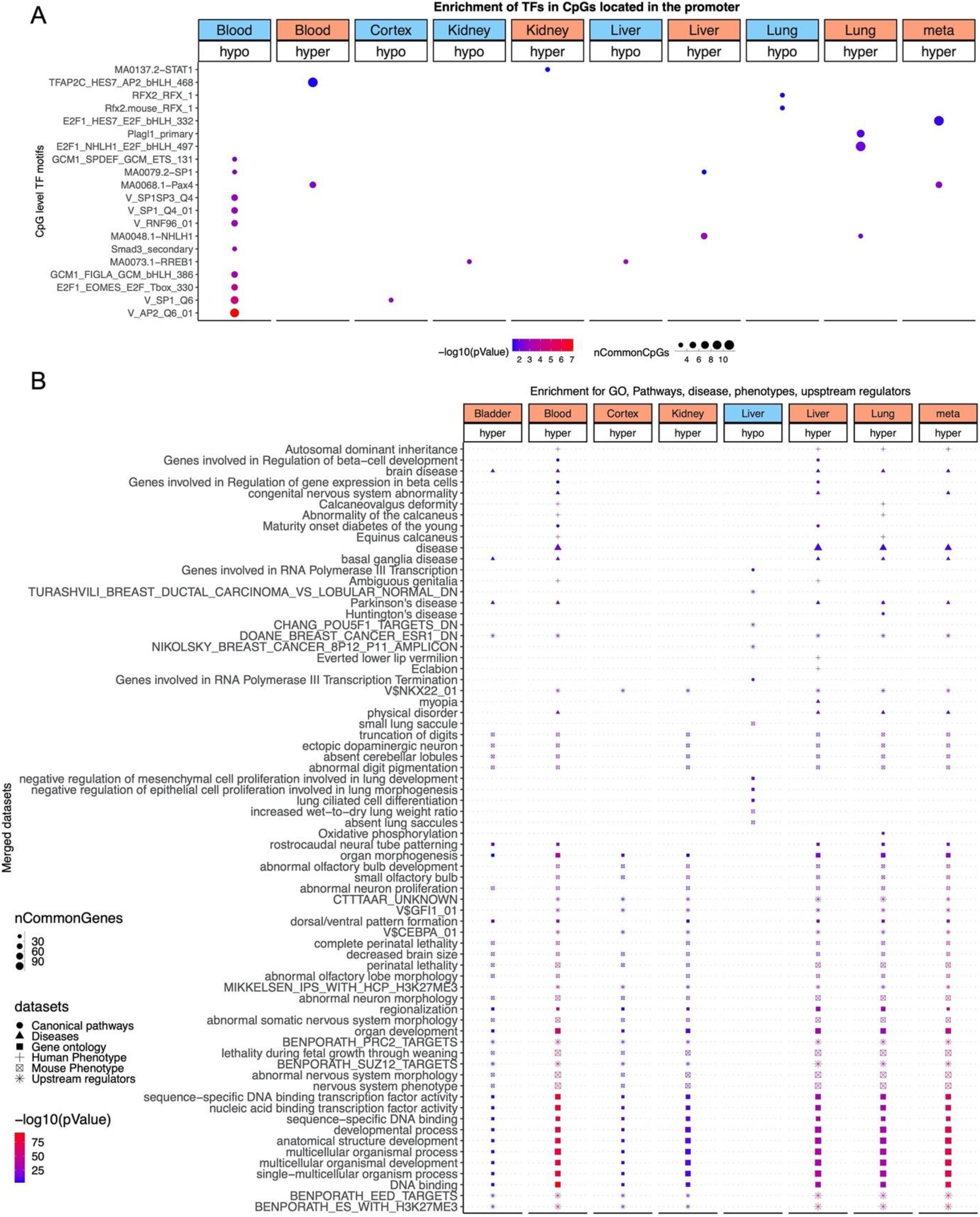
Enrichment analysis of the top DNAm aging marks in porcine tissues. A) Transcriptional motif enrichment for the top CpGs in the promoter (and 5 prime UTR) of the neighboring genes. The motifs were scanned using the FIMO for all the probes, and the enrichment was tested using a hypergeometric test. B) Gene set enrichment analysis of proximate genes with DNAm aging in porcine tissues. The analysis was done using the genomic region of enrichment annotation tool ^56^. The gene level enrichment was done using GREAT analysis ^56^ with the human Hg19 background. The background probes were limited to 23,087 probes that were mapped to the same gene in the pig genome. The top three enriched datasets from each category (Canonical pathways, diseases, gene ontology, human and mouse phenotypes, and upstream regulators) were selected and further filtered for significance at p < 10^-8^.

While identification of genes proximal to age-associated CpGs, as performed above, are useful, the likelihood of their potential effects on cells is difficult to gauge. This can be partly addressed by carrying out analysis to identify enrichment of implicated genes in specific pathways, pathologies and biological processes (**Figure 3**). This analysis highlighted the following features impacted by age-related CpG methylation changes in porcine cells: organism development, the nervous system and metabolism; all of which have also been found to be implicated in epigenetic aging of humans and other species (**Figure 3B**). Furthermore, genes proximal to hypermethylated CpGs are associated with H3K27Me3 marks and are often polycomb protein EED targets in porcine tissues. EED is a member of the multimeric Polycomb family protein complex that maintains the transcriptional repressive states of genes. These proteins also regulate H3K27Me3 marks, DNA damage, and senescence states of the cells during aging ^38^.

### Studying differences across pig lines

With the DNA methylation data sets we have accrued from the different porcine breeds, we are in a position to identify CpGs that differ between domestic and minipig breeds. We compared DNA methylation profiles from blood of domestic pigs with those from Wisconsin Miniature Swine™. We found found the mean methylation across CpGs located in CpG islands is higher in minpigs than in the other two pig lines (**Supplementary Figure 4**). Similarly, the average rate of change in methylation across island CpGs is increased in minipigs (**Supplementary Figure 4**).

We analyzed individual CpGs using two different multivariate models. In the first, the DNAm levels of a given CpG were regressed on age and breed (minipig versus reset) to identify CpGs that are associated with aging in both breeds (age main effect). This model was also used to identify CpGs with significantly different basal methylation levels between breeds (Minipig main effect). The second model identified age-related CpGs with different rates of methylation changes between these two pig breeds independent of the direction of change. At a genome-wide significance set at p < 1e-8; 10,167 age-associated CpGs were shared between both pig breeds, while 825 CpGs had different baseline methylation levels, and the rate of methylation change of 32 age-associated CpGs were significantly different between the two (**Figure 4A**). Thus, while age has the largest impact on methylation of these CpGs, there is an inherent species-specific difference in basal methylation levels of a substantial number of CpGs, which is expected given the overt differences between the breeds. The top CpGs with divergent rate of methylation change between the breeds are proximal to the *MGST1* exon, *SON* 3’UTR, and *TFAP2B* exon **(Figure 4A-C**). CpGs with different basal methylation levels between the two breeds were located within the *OSBP* exon, *ENC1* intron, and *CTNNBL1* upstream regions (**Figure 4A**). In total, eight categories of CpGs can be defined based on the direction of their methylation change with age in the two pig breeds. While methylation of most age-associated CpGs was altered in similar directions in both breeds (hypo or hyper in both axes), some were clearly in opposite directions (hyper in one axis and hypo in the other). The *LMNA* intron is an example region that displayed extreme divergence. While its intron was hypomethylated with age in the domestic pig, it was hypermethylated in minipigs (**Figure 4C**). An enrichment analysis of age-related CpGs that are exclusive to either breed implicated pathways involved in development, survival, cancer, and growth (**Figure 4D**).

**Figure 4.**
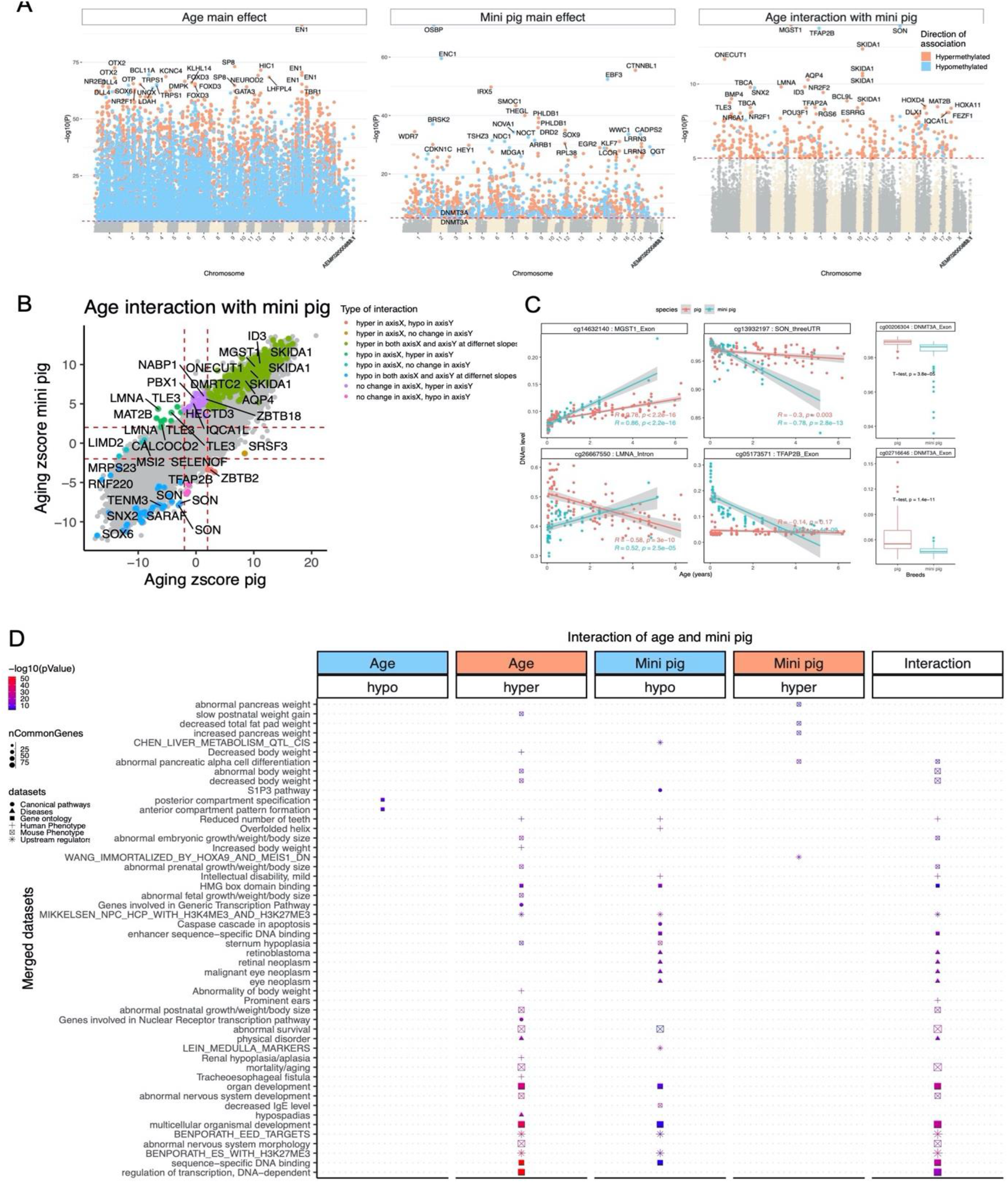
EWAS of multivariate regression models for pig species and age. Aging differences between domestic and minipig breeds. The figure visualizes the results of 2 different linear regression models which used individual CpGs as dependent variable. The 2 linear models differ in terms of the underlying covariates: two covariates (age and pig breed) or three covariates (age, pig breed, and interaction effect). A) Manhattan plots of DNAm aging loci that are shared between minipigs and domestic pigs (Aging main effect), basal breed differences in DNAm levels (minipig main effect), and the interaction of breed and aging, which represent the loci with a divergent DNAm aging pattern between minipigs and domestic pigs. The analysis is done by multivariate regression models with or without (to estimate the main effect) interaction term for age and breeds. For breeds, the domestic pig is the reference variable to estimate the direction of change. Sample sizes: domestic pigs, 98; minipigs, 60. The red line in the Manhattan plot indicates p <1E-35. B) Scatter plots of DNAm aging between minipigs and domestic pigs. The highlighted CpGs are the loci with significant DNAm aging interaction between breeds at a 5% FDR rate. In total, eight categories of interaction were defined based on the aging Z-statistic of each breed. C) Age versus methylation levels for select CpGs with significant interaction terms between breed and age. D) Enrichment analysis of the genes proximate to CpGs related to age (shared between breeds), minipig, and age:minipig interaction. The gene-level enrichment was done using GREAT analysis ^56^ and human Hg19 background. The background probes were limited to 23,087 probes that were mapped to the same gene in the pig genome. The top CpGs were selected at a p<1E-5 and based on Beta values of association for up to 500 in a positive or negative sign.

Our analysis of differentially methylated CpGs between domestic and minipigs implicated genes that regulate weight. This particularly interesting finding prompted us to query whether any of these identified genes are associated with weight differences in humans. To this end, we overlapped the EWAS genes with a large GWAS meta-analysis of body-mass index (BMI) in humans, which included 681,275 participants in UK Biobank and GIANTBMI consortium ^39^. Strikingly, several of the EWAS-identified genes had genetic variants that are associated with increased BMI in humans (**Figure 5**). The top genes include *ADCY3, TFAP2B, SKOR1*, and *GPR61*, which had numerous SNPs in different gene regions associated with human BMI. Thus, our EWAS analysis of pig breeds of overtly different sizes implicates genes that were implicated by a published GWAS of human BMI.

**Figure 5.**
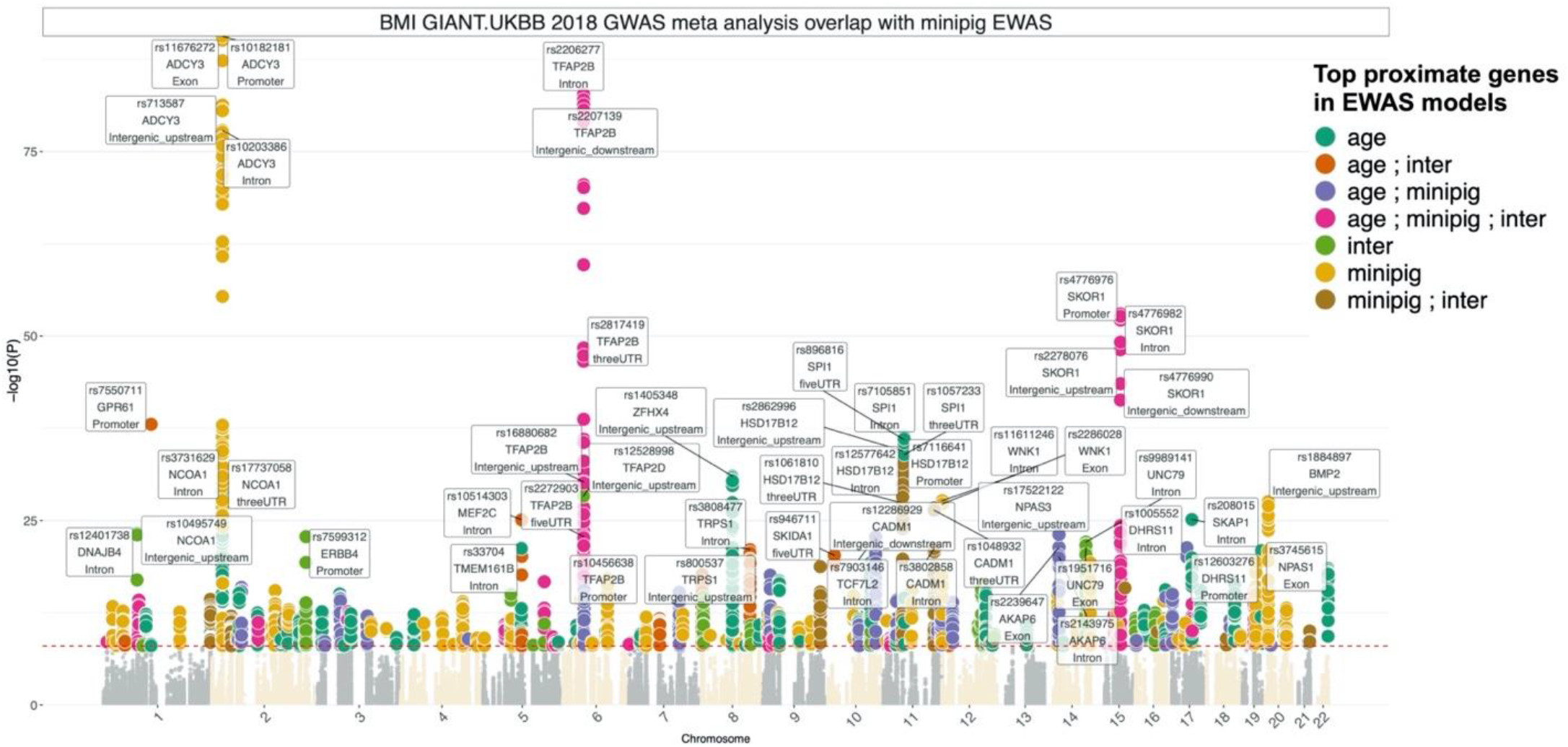
Overlapping EWAS results in pigs with GWAS results in humans. EWAS of aging, minipig, and aging:pig breed interaction identifies genes with genetic variants associated with *human* body mass index. The GWAS is based on summary statistics of BMI meta-analysis of 681,275 participants in the UK Biobank and GIANT_BMI_ consortium ^39^. The coloring is based on the genes identified by one or multiple EWAS of age, minipig, and age:minipig interaction (inter). The labels indicate the top SNP from each of the top 30 gene regions.

## Discussion

We have previously developed several human epigenetic clocks from DNA methylation profiles that were derived from various versions of human Illumina DNA methylation arrays. As these arrays are specific to the human genome, a critical step toward crossing the species barrier was the development and use of a mammalian DNA methylation array that profiles 36 thousand CpGs with flanking DNA sequence that are highly conserved across numerous mammalian species. The employment of this array to profile 238 porcine tissues from 3 pig lines, represents the most comprehensive epigenetic dataset of domestic pigs thus far. These data allowed us to construct four highly accurate DNA methylation-based age estimators for three pig lines. Two of these clocks apply only to pigs: the pan-tissue clock which was trained on methylation profiles of six tissues, is expected to apply to most pig tissues. The blood clock on the other hand, was trained using only blood DNA methylation data. These two highly-accurate porcine clocks are readily and easily included in porcine-based models of diseases and health conditions. This will encourage investigations into the relationship between age and diseases, and also uncover the effects of environment, living condition, food, medicine and treatment on the rate of porcine epigenetic aging. While toxicity, mutagenicity and carcinogenicity are considered health-impacting effects that draw the attention of health experts, age-accelerants have yet to be appreciated as being potentially as important, or perhaps even more so, as age is the biggest risk factor for almost all diseases, and accelerated epigenetic aging is associated with a whole host of pathologies and health conditions. While mice can be used to detect potential age-accelerating agents, this route is evidently limited due to the inherent differences between humans and mice, such as food, sleep-wake cycle, metabolism, physiology, endocrinology, and disease susceptibility. The better compatibility between pigs and humans are elaborated above, and the availability of these porcine epigenetic clocks further consolidates the greater relevance of employing porcine models for human health conditions. This advantage is compounded by the successful generation of human-pig clocks that apply to both species. It is essential to appreciate the profound significance of these dual species clocks. With a single mathematical formula, the age of humans and pigs can be estimated based solely on methylation levels of a few hundred cytosines. At the most fundamental level, this demonstrates that the mechanisms and processes that underpin aging in these two mammalian species that diverged millions of years ago are essential for life. Species diversification and adaptation through selection would gain specialized and advantageous features and lose others, but features essential for life will be retained. As such, the mechanisms that underlie aging must be essential for life as well. This is consistent with the fact that some of the loci that harbor age-associated CpGs in pigs and humans (as well as other mammalian species), are bivalent chromatin domains and targets of the polycomb repressive complex (PRC). This is of significance because these regions are primarily located upstream of Hox genes which specify development of various parts of the body of organisms ranging from worms, flies, mice and humans. In other words, these are some of the oldest genetic elements of life. In this study the highest scoring porcine blood CpG is one that resides in the promoter if the engrailed gene (EN1), which is a member of the Hox gene family. EN1 also scored the highest in meta-analyses of all the tissues. It is as yet unclear how methylation of these loci are involved with aging, but their increased methylation with age hints at the possible role of cellular identity and differentiation. This is no doubt an exciting avenue of exploration in which these epigenetic clocks will be essential.

The considerable similarities between humans and pigs allows for the testing of age-mitigation interventions as well as factors that impact longevity in pigs, which have a shorter lifespan and are much more amenable subjects for controlled trials. To accurately translate age-related findings from pigs to humans however requires a correct and accurate measure of age-equivalence. We fulfilled this need through a two-step process. The first, which we described above is the generation of dual-species clocks (human-pig), one of which is as accurate in estimating pig age as it is for human age; in chronological units of years. The second is the expression of pig and human ages in respect to the maximum recorded ages of their respective species (species lifespan), i.e. 23 years for pigs and 122 years for humans. The mathematical operation of generating a ratio eliminates chronological units of time and produces a value that indicates the age of the organism in respect to the maximum age of its own species. This allows a meaningful and realistic cross-species comparison of biological age. For example, the biological fitness of a 20 year-old pig, which is very old, is not equivalent to that of a 20 year-old human, who is young. However, a pig with a relative epigenetic age of 0.5 is more comparable to a human of similar relative epigenetic age. Collectively, the ability to use a single mathematical formula to measure epigenetic age of different species and the replacement of chronological units of time with proportion of lifespan, are two significant innovations that will propel cross-species research as well as cross-species benefits.

In addition to age-related epigenetic changes, we also compared DNA methylation profiles between domestic and minipigs. CpGs with substantially different basal methylation levels between the two breeds were identified proximal to the *OSBP* gene. This is of great significance, as OSBP is a transport protein that translocates sterols from lysosomes into the nucleus where the sterol represses the expression of the *LDL* receptor gene, *HMG-CoA* reductase gene and *HMG* synthase gene. The reduction of LDL receptor increases the risk of atherosclerosis and cardiovascular disease. Indeed, mice with ablated *LDL* receptor genes develop plaques in their aortas, while wildtype mice are free of such plaques. Compounding the effects on LDL receptor levels is the effect on expression of HMG-CoA and HMG synthase, which are members of the intracellular pathway that synthesize cholesterol. These features are of particular interest given the fact that while domestic pigs are refractive to atherosclerotic plaque development, minipigs are susceptible and hence used as models for cardiovascular disease. Another feature of interest is the identification of differentially methylated CpGs associated with genes involved in the development of obesity. The matching of EWAS targets with a large GWAS meta-analysis of body-mass index (BMI) in humans led to the identification of overlapping genes included *TFAP2B*, which influences the effect of dietary fat on weight ^40^, *GPR61*, which is involved in the regulation of food intake and body weight ^41,42^, and *ADCY3* and *SKOR1*, both of which are associated with obesity and BMI ^43–46^. Collectively, these associations points to the contribution of epigenetic control and influence on weight gain and obesity, in addition to highlighting the translational relevance of porcine models for cardiovascular disease and obesity-related research. It is interesting to note that the *Dnmt3a* locus is also differentially methylated between these two breeds. It is tempting to speculate that this may be an upstream event that impacts on the downstream methylation differences described above.

Our study is limited in that all animals were younger than 6.3 years old while the maximum observed lifespan of domestic pigs (*Sus scrofa domesticus*) appears to be 23. In our own data base, we assigned the Wisconsin Miniature Swine™ the Latin name “*Sus scrofa minusculus”* with an estimated maximum lifespan of 23 years.

As porcine biomedical models for a wide range of age-related disorders are currently in use or being developed, including Alzheimer’s disease ^47^, cardiovascular disease ^48^, diabetes ^22^, and cancer models ^21,23,24^, the availability of these epigenetic clocks will extend the use of porcine models for aging, and possibly obesity studies.

## Materials and Methods

### Porcine samples

All animal procedures were approved by the University of Illinois and University of Wisconsin Institutional Animal Care and Use Committee, and all animals received humane care according to the criteria outlined in the Guide for the Care and Use of Laboratory Animals. Porcine whole blood samples (n=146) were collected from female Large White X Landrace crossbred domestic pigs (n=84, age range 11 – 2,285 days) and Wisconsin Miniature Swine™ (n=60, age range 8 – 1,880 days) at the University of Wisconsin-Madison. Whole blood (n=16) and tissue samples (bladder, frontal cortex, kidney, liver, lung; n=16/tissue type) were collected from 16 Large White X Minnesota minipig crossbred pigs (n=9 female, n=8 male, age range 29 – 1,447 days) at the University of Illinois at Urbana-Champaign. All blood samples were collected in EDTA tubes, aliquoted, and flash frozen in liquid nitrogen within 10 minutes of collection. Tissue samples were collected and flash frozen within 10 minutes of euthanasia. All samples were stored at −80 until processing. Samples were shipped to the University of California, Los Angeles Technology Center for Genomics & Bioinformatics for DNA extraction and generation of DNA methylation data.

### Human tissue samples

To build the human-pig clock, we analyzed previously generated methylation data from n=850 human tissue samples (adipose, blood, bone marrow, dermis, epidermis, heart, keratinocytes, fibroblasts, kidney, liver, lung, lymph node, muscle, pituitary, skin, spleen) from individuals whose ages ranged from 0 to 93. The tissue samples came from three sources. Tissue and organ samples from the National NeuroAIDS Tissue Consortium ^49^. Blood samples from the Cape Town Adolescent Antiretroviral Cohort study ^50^. Skin and other primary cells provided by Kenneth Raj ^51^. Ethics approval (IRB#15-001454, IRB#16-000471, IRB#18-000315, IRB#16-002028).

### DNA methylation data

We generated DNA methylation data using the custom Illumina chip “HorvathMammalMethylChip40”. The mammalian methylation array provides high sequencing depth (over thousand X) of highly conserved CpGs in mammals, but focuses only on 36k CpGs that are highly conserved across mammals. Out of 38,000 probes on the array, 2,000 were selected based on their utility for human biomarker studies: these CpGs, which were previously implemented in human Illumina Infinium arrays (EPIC, 450K) were selected due to their relevance for estimating age, blood cell counts, or the proportion of neurons in brain tissue. The remaining 35,988 probes were chosen to assess cytosine DNA methylation levels in mammalian species (Arneson, Ernst, Horvath, in preparation). The particular subset of species for each probe is provided in the chip manifest file at the NCBI Gene Expression Omnibus (GEO) platform (GPL28271). The SeSaMe normalization method was used to define beta values for each probe ^52^.

### Relative age estimation

To introduce biological meaning into age estimates of pigs and humans that have very different lifespan; as well as to overcome the inevitable skewing due to unequal distribution of data points from pigs and humans across age range, relative age estimation was made using the formula: Relative age= Age/maxLifespan where the maximum lifespan for the two species was chosen from an updated version of the *anAge* data base ^53^.

### Clocks and penalized regression

Details on the clocks (CpGs, genome coordinates) and R software code are provided in the Supplement. Penalized regression models were created with glmnet ^54^. We investigated models produced by elastic net” regression (alpha=0.5 in glmnet R function). The optimal penalty parameters in all cases were determined automatically by using a 10 fold internal cross-validation (cv.glmnet) on the training set. By definition, the alpha value for the elastic net regression was set to 0.5 (midpoint between Ridge and Lasso type regression) and was not optimized for model performance. We performed a cross-validation scheme for arriving at unbiased (or at least less biased) estimates of the accuracy of the different DNAm based age estimators. One type consisted of leaving out a single sample (LOOCV) from the regression, predicting an age for that sample, and iterating over all samples. We subset the set of CpG probes to those that uniquely mapped to a CpG site in the swine genome. While no transformation was used for the blood clock for pigs, we did use a log linear transformation for the dual species clock of chronological age (Supplement). The accuracy of the resulting clocks was assessed via cross validation estimates of 1) the correlation R between the predicted epigenetic age and the actual (chronological) age of the animal, 2) the median absolute difference between DNAm age and chronological age (mae).

### EWAS and Functional Enrichment

EWAS was performed in each tissue separately (bladder, blood, cerebral cortex, kidney, liver, lung) using the R function “standardScreeningNumericTrait” from the “WGCNA” R package^55^. Next the results were combined across tissues using Stouffer’s meta-analysis method. The functional enrichment analysis was done using the genomic region of enrichment annotation tool ^56^. CpGs implicated by our EWAS were filtered for CpG position information, lifted over to the human genome using UCSC’s Liftover tool and fed into the online functional analysis tool GREAT using the default mode, to obtain a list of significantly enriched functions for both positive and negative EWAS hits in the different tissues.

### Transcription factor enrichment and chromatin states

The FIMO (Find Individual Motif Occurrences) program scans a set of sequences for matches of known motifs, treating each motif independently ^57^. We ran TF motif (FIMO) scans of all probes on the HorvathMammalMethyl40 chip using motif models from TRANSFAC, UniPROBE, Taipale, Taipaledimer and JASPAR databases. A FIMO scan p-value of 1E-4 was chosen as the cutoff (lower FIMO p-values reflect a higher probability for the local DNA sequence matching a given TF motif model). This cutoff implies that we would find almost all TF motif matches that could possibly be associated with each site, resulting in an abundance of TF motif matches. We caution the reader that our hypergeometric test enrichment analysis did not adjust for CG content.

## Author contributions

SH conceived of the study and wrote the article. The remaining authors helped with the statistical analysis or the data generation. All authors reviewed and edited the article.

## Acknowledgements

This work was supported by the Paul G. Allen Frontiers Group (SH). We would like to thank Dr. Ana Cecilia Escobar Lopez, Jamie Reichert, Jennifer Frank, Justin Hickman, Keri Graff and Nathan Chesmore for their technical assistance during the sampling of the Wisconsin Miniature Swine.

## Supplementary Material

**Supplementary Figure 1.**
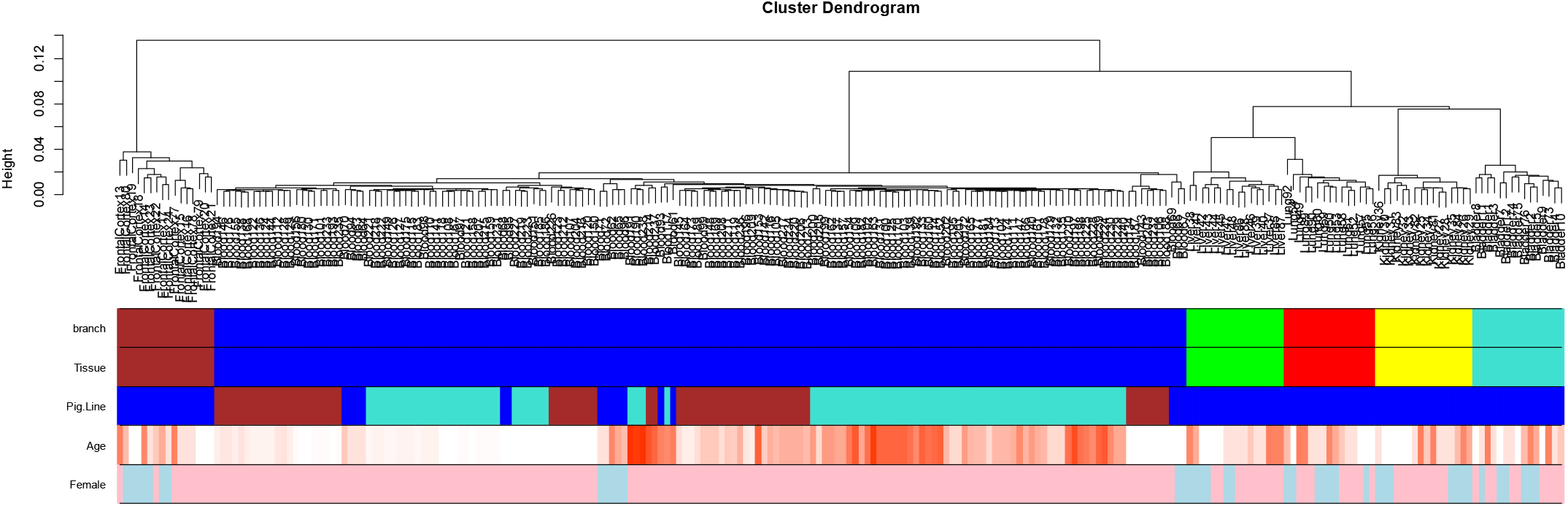
Unsupervised hierarchical clustering of porcine tissues. Average linkage hierarchical clustering based on the interarray correlation coefficient (Pearson correlation). Contrasting the first color band (based on cluster branches) with the second color band shows that the arrays cluster by tissue type (blood=blue, bladder=turquoise, frontal brain cortex=brown, kidney=yellow, liver=green, lung=red). Pig line: turquoise=domestic, brown=Wisconsin Miniature Swine, blue= Domestic/Minnesota Mini Cross. The last color encodes sex (pink=female).

**Supplementary Figure 2.**
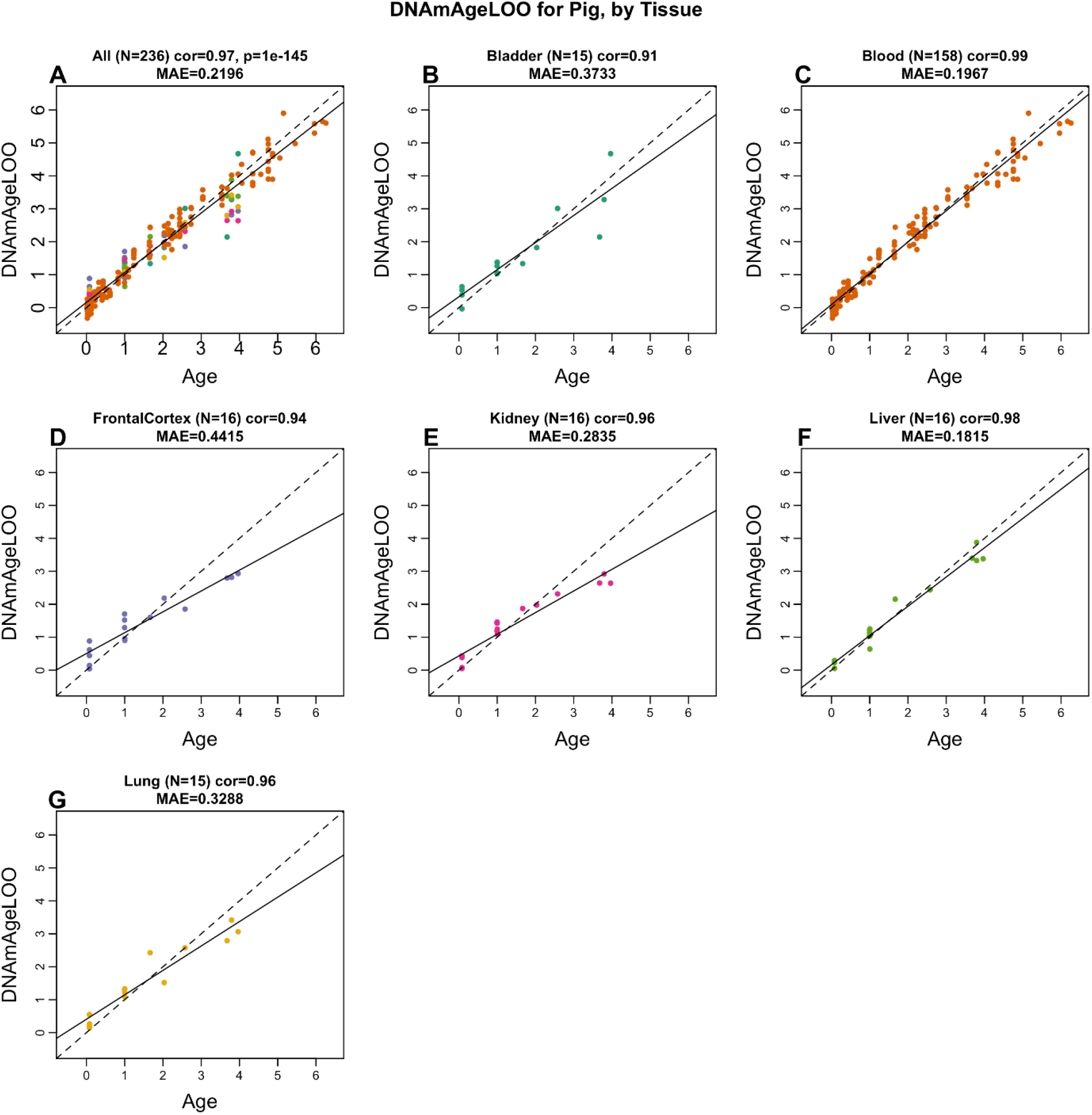
Pan tissue clock for pigs applied to individual tissues. A) All porcine tissues combined. B) Bladder, C) Blood, D) Frontal cortex (brain), E) Kidney, F) Liver, G) Lung. Each panel reports the leave-one-out estimate of age (y-axis) versus chronological age (in years). Each title reports the sample size, Pearson correlation coefficient, and median absolute error.

**Supplementary Figure 3.**
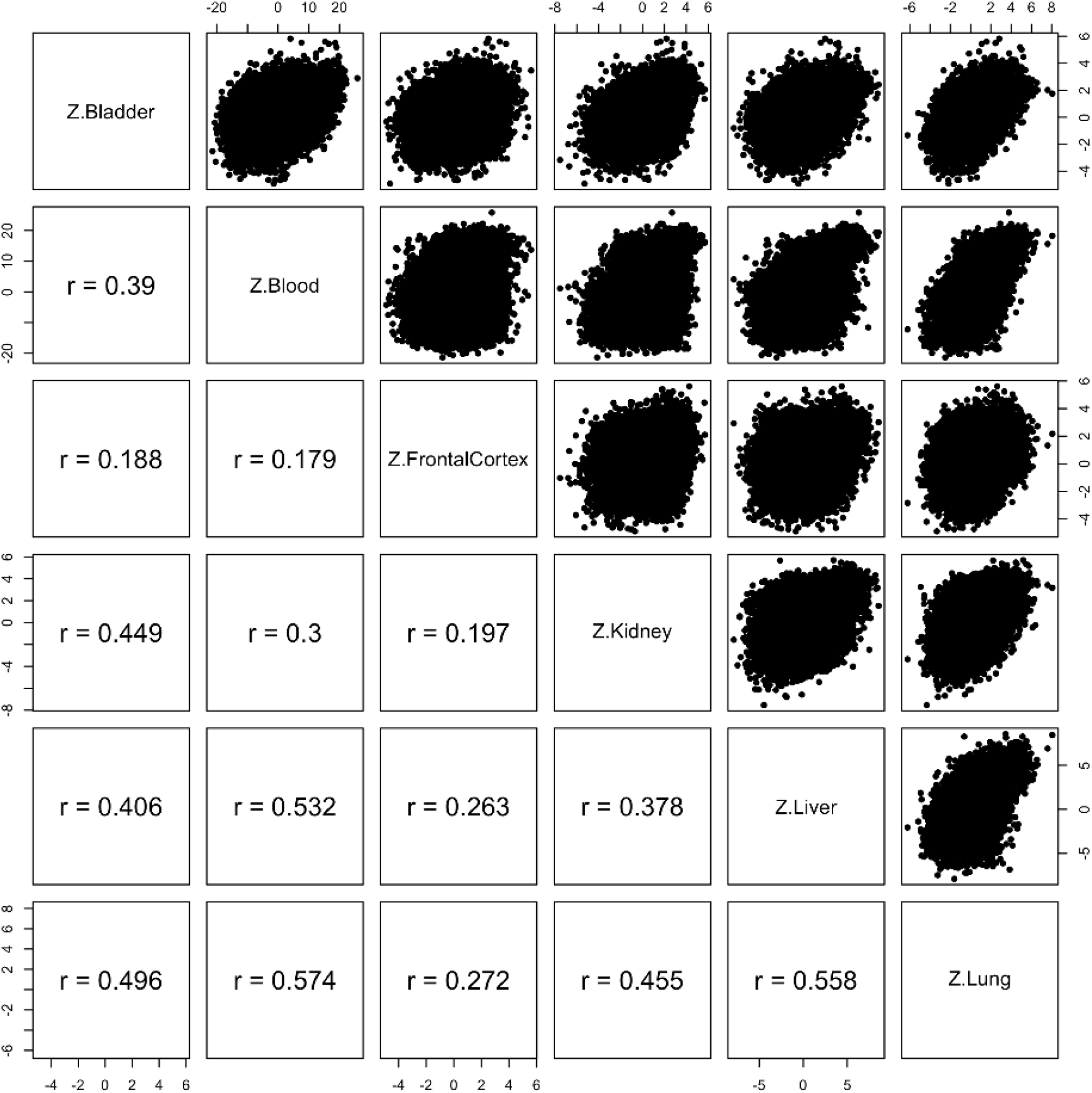
Epigenome wide association study of correlation in different tissues. Each dot corresponds to a CpG. Z statistics for a correlation test of age in the respective tissues.

**Supplementary Figure 4.**
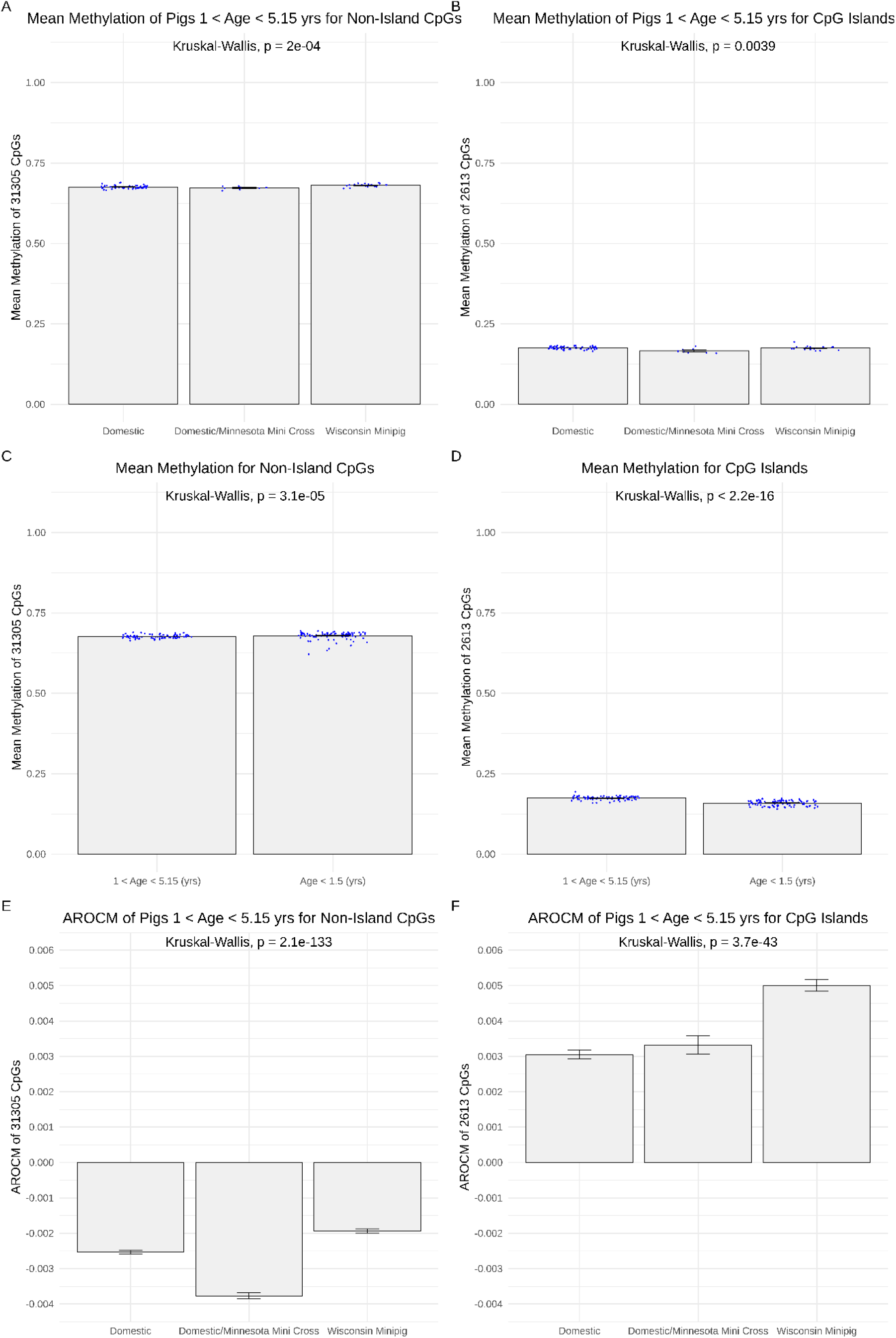
Mean Methylation and Average Rate of Change of Methylation. A, B, C, D) Mean methylation of all CpG sites that (A, C) map to the Sus scrofa genome on a custom mammalian methylation array and (B, D) occur only within CpG islands. Figures (A) and (B) contain samples between the ages of 1 and 5.15 years old and are grouped by breed. Figures (C) and (D) group samples into two age ranges: between 1 and 5.15 years old, and less than 1.5 years old. Blood samples were taken from three breeds: “Domestic” pigs, a cross between “Domestic” and “Minnesota Mini” pigs, and “Wisconsin Minipigs”. Each point represents the mean methylation of all selected CpG sites in a single sample. The standard error was calculated using the standard deviation of the average methylation across the individual samples. E,F) Average rate of change of methylation (AROCM) of all CpG sites that (E) map to the Sus scrofa genome on a custom mammalian methylation array and (F) occur only within CpG islands. Figures (E) and (F) contain samples between the ages of 1 and 5.15 years old and are grouped by breedBlood samples were taken from three breeds: “Domestic” pigs, a cross between “Domestic” and “Minnesota Mini” pigs, and “Wisconsin Minipigs”. AROCM is defined for each breed as the average of the rates of change at the selected CpG sites. The rate of change at each CpG is the slope coefficient of a simple linear regression where the beta values of a single CpG are regressed on the ages of the samples. The standard error was calculated using the standard deviation of the rate of change of methylation across the selected CpG sites. To ensure a balanced statistical design, we restricted the analysis to pigs aged between 1 and 5.2. The title of each figure reports a non-parametric group comparison test (Kruskal Wallis tests).

## References

1. Kumar, S. & Hedges, S.B. A molecular timescale for vertebrate evolution. Nature 392, 917–920 (1998).

2. Meredith, R.W. et al. Impacts of the Cretaceous Terrestrial Revolution and KPg extinction on mammal diversification. Science 334, 521–4 (2011).

3. Caliebe, A., Nebel, A., Makarewicz, C., Krawczak, M. & Krause-Kyora, B. Insights into early pig domestication provided by ancient DNA analysis. Sci Rep 7, 44550 (2017).

4. Groenen, M.A. et al. Analyses of pig genomes provide insight into porcine demography and evolution. Nature 491, 393–8 (2012).

5. Swindle, M.M., Makin, A., Herron, A.J., Clubb, F.J., Jr. & Frazier, K.S. Swine as models in biomedical research and toxicology testing. Vet Pathol 49, 344–56 (2012).

6. Schook, L.B. et al. DNA-based animal models of human disease: from genotype to phenotype. Dev Biol (Basel) 132, 15–25 (2008).

7. Schachtschneider, K.M. et al. The Oncopig Cancer Model: An Innovative Large Animal Translational Oncology Platform. Front Oncol 7, 190 (2017).

8. Schachtschneider, K.M. et al. Adult porcine genome-wide DNA methylation patterns support pigs as a biomedical model. BMC Genomics 16, 743 (2015).

9. Choi, M. et al. Genome-wide analysis of DNA methylation in pigs using reduced representation bisulfite sequencing. DNA Res 22, 343–55 (2015).

10. Schook, L.B. et al. Unraveling the swine genome: implications for human health. (2015).

11. Gutierrez, K., Dicks, N., Glanzner, W.G., Agellon, L.B. & Bordignon, V. Efficacy of the porcine species in biomedical research. Frontiers in genetics 6, 293 (2015).

12. Santulli, G. et al. Models for preclinical studies in aging-related disorders: One is not for all. Translational medicine @ UniSa 13, 4–12 (2016).

13. Ganderup, N.C., Harvey, W., Mortensen, J.T. & Harrouk, W. The minipig as nonrodent species in toxicology--where are we now? Vol. 31 507–528 (2012).

14. Heino, T.J., Alm, J.J., Moritz, N. & Aro, H.T. Comparison of the osteogenic capacity of minipig and human bone marrow-derived mesenchymal stem cells. Journal of Orthopaedic Research 30, 1019–1025 (2012).

15. Schwartz, R.S. et al. Drug-eluting stents in preclinical studies: updated consensus recommendations for preclinical evaluation. Circulation: Cardiovascular Interventions 1, 143–153 (2008).

16. Coronel, R. et al. Dietary n-3 fatty acids promote arrhythmias during acute regional myocardial ischemia in isolated pig hearts. Cardiovascular research 73, 386–394 (2007).

17. Dixon, J.A. & Spinale, F.G. Large animal models of heart failure: a critical link in the translation of basic science to clinical practice. Circulation: Heart Failure 2, 262–271 (2009).

18. Ekeløf, S., Rosenberg, J., Jensen, J.S. & Gögenur, I. Pharmacological attenuation of myocardial reperfusion injury in a closed-chest porcine model: a systematic review. Journal of cardiovascular translational research 7, 570–580 (2014).

19. Al-Mashhadi, R.H. et al. Familial hypercholesterolemia and atherosclerosis in cloned minipigs created by DNA transposition of a human PCSK9 gain-of-function mutant. Science translational medicine 5, 166ra1–166ra1 (2013).

20. Shim, J., Al-Mashhadi, R.H., Sørensen, C.B. & Bentzon, J.F. Large animal models of atherosclerosis-new tools for persistent problems in cardiovascular medicine. The Journal of Pathology 238, 257–266 (2016).

21. Schook, L.B. et al. A Genetic Porcine Model of Cancer. in PloS one Vol. 10 e0128864 (2015).

22. Wolf, E., Braun-Reichhart, C., Streckel, E. & Renner, S. Genetically engineered pig models for diabetes research. Transgenic research 23, 27–38 (2014).

23. Gaba, R.C. et al. Development and comprehensive characterization of porcine hepatocellular carcinoma for translational liver cancer investigation. Oncotarget 11, 2686–2701 (2020).

24. Kalla, D., Kind, A. & Schnieke, A. Genetically Engineered Pigs to Study Cancer. Int J Mol Sci 21(2020).

25. Al-Mashhadi, R.H. et al. Diabetes with poor glycaemic control does not promote atherosclerosis in genetically modified hypercholesterolaemic minipigs. Diabetologia 58, 1926–1936 (2015).

26. Elmadhun, N.Y., Lassaletta, A.D., Chu, L.M. & Sellke, F.W. Metformin alters the insulin signaling pathway in ischemic cardiac tissue in a swine model of metabolic syndrome. The Journal of thoracic and cardiovascular surgery 145, 258–266 (2013).

27. Schachtschneider, K.M. et al. A validated, transitional and translational porcine model of hepatocellular carcinoma. Oncotarget 8, 63620–63634 (2017).

28. Horvath, S. DNA methylation age of human tissues and cell types. Genome Biol 14, R115 (2013).

29. Horvath, S. et al. Accelerated epigenetic aging in Down syndrome. Aging Cell 14, 491–5 (2015).

30. Horvath, S. & Levine, A.J. HIV-1 Infection Accelerates Age According to the Epigenetic Clock. J Infect Dis 212, 1563–73 (2015).

31. Horvath, S. et al. Obesity accelerates epigenetic aging of human liver. Proc Natl Acad Sci U S A 111, 15538–43 (2014).

32. Horvath, S. & Raj, K. DNA methylation-based biomarkers and the epigenetic clock theory of ageing. Nat Rev Genet (2018).

33. Marioni, R. et al. DNA methylation age of blood predicts all-cause mortality in later life. Genome Biol. 16, 25 (2015).

34. Marioni, R.E. et al. The epigenetic clock is correlated with physical and cognitive fitness in the Lothian Birth Cohort 1936. Int J Epidemiol 44, 1388–96 (2015).

35. Chen, B.H. et al. DNA methylation-based measures of biological age: meta-analysis predicting time to death. Aging (Albany NY) 8, 1844–1865 (2016).

36. Lu, A.T. et al. DNA methylation GrimAge strongly predicts lifespan and healthspan. Aging (Albany NY) 11, 303–327 (2019).

37. Wege, H. et al. Regeneration in pig livers by compensatory hyperplasia induces high levels of telomerase activity. Comp Hepatol 6, 6 (2007).

38. Ito, T., Teo, Y.V., Evans, S.A., Neretti, N. & Sedivy, J.M. Regulation of Cellular Senescence by Polycomb Chromatin Modifiers through Distinct DNA Damage- and Histone Methylation-Dependent Pathways. Cell Rep 22, 3480–3492 (2018).

39. Yengo, L. et al. Meta-analysis of genome-wide association studies for height and body mass index in approximately 700000 individuals of European ancestry. Hum Mol Genet 27, 3641–3649 (2018).

40. Stocks, T. et al. TFAP2B influences the effect of dietary fat on weight loss under energy restriction. PLoS One 7, e43212 (2012).

41. Nambu, H. et al. Characterization of metabolic phenotypes of mice lacking GPR61, an orphan G-protein coupled receptor. Life Sci 89, 765–72 (2011).

42. Felix, J.F. et al. Genome-wide association analysis identifies three new susceptibility loci for childhood body mass index. Hum Mol Genet 25, 389–403 (2016).

43. Goni, L. et al. Interaction between an ADCY3 Genetic Variant and Two Weight-Lowering Diets Affecting Body Fatness and Body Composition Outcomes Depending on Macronutrient Distribution: A Randomized Trial. Nutrients 10(2018).

44. Grarup, N. et al. Loss-of-function variants in ADCY3 increase risk of obesity and type 2 diabetes. Nat Genet 50, 172–174 (2018).

45. Saeed, S. et al. Loss-of-function mutations in ADCY3 cause monogenic severe obesity. Nat Genet 50, 175–179 (2018).

46. Kaewsutthi, S. et al. Exome sequencing in Thai patients with familial obesity. Genet Mol Res 15(2016).

47. Hoffe, B. & Holahan, M.R. The Use of Pigs as a Translational Model for Studying Neurodegenerative Diseases. Front Physiol 10, 838 (2019).

48. Crisóstomo, V. et al. Common swine models of cardiovascular disease for research and training. Lab Anim (NY) 45, 67–74 (2016).

49. Morgello, S. et al. The National NeuroAIDS Tissue Consortium: a new paradigm in brain banking with an emphasis on infectious disease. Neuropathol Appl Neurobiol 27, 326–35. (2001).

50. Horvath, S. et al. Perinatally acquired HIV infection accelerates epigenetic aging in South African adolescents. AIDS (London, England) 32, 1465–1474 (2018).

51. Kabacik, S., Horvath, S., Cohen, H. & Raj, K. Epigenetic ageing is distinct from senescence-mediated ageing and is not prevented by telomerase expression. Aging (Albany NY) 10, 2800–2815 (2018).

52. Zhou, W., Triche, T.J., Jr, Laird, P.W. & Shen, H. SeSAMe: reducing artifactual detection of DNA methylation by Infinium BeadChips in genomic deletions. Nucleic Acids Research 46, e123–e123 (2018).

53. de Magalhaes, J.P., Costa, J. & Church, G.M. An analysis of the relationship between metabolism, developmental schedules, and longevity using phylogenetic independent contrasts. J Gerontol A Biol Sci Med Sci 62, 149–60 (2007).

54. Friedman, J., Hastie, T. & Tibshirani, R. Regularization Paths for Generalized Linear Models via Coordinate Descent. Journal of Statistical Software 33, 1–22 (2010).

55. Langfelder, P. & Horvath, S. WGCNA: an R package for weighted correlation network analysis. BMCBioinformatics 9, 559 (2008).

56. McLean, C.Y. et al. GREAT improves functional interpretation of cis-regulatory regions. Nat Biotechnol 28(2010).

57. Bailey, T.L. et al. MEME Suite: tools for motif discovery and searching. Nucleic Acids Research 37, W202–W208 (2009).

